# The phosphatidylserine-binding proteins Turandots protect the peripheral nervous system from antimicrobial peptide toxicity

**DOI:** 10.64898/2026.04.30.721952

**Authors:** Samuel Rommelaere, Siqi Wang, Samuel Vernon, Kenan Krakovic, Jean-Philippe Boquete, Brian D. McCabe, Bruno Lemaitre

## Abstract

Inflammation increases with aging and contributes to neurodegeneration, yet the principles that determine how immune effectors target host neural tissue remain poorly understood. Antimicrobial peptides (AMPs) are central components of innate immune defenses strongly induced upon infection and upon aging. Studies have shown that AMPs can exhibit cytotoxicity toward host cells, pointing to a role in neurodegeneration. We show that cationic AMPs selectively bind and damage motoneurons that expose phosphatidylserine (PS), an anionic phospholipid normally restricted to the inner leaflet of the plasma membrane. Both infection and aging increase neuronal PS exposure alongside AMP expression. AMP binding occurs in a PS-dependent manner, leading to synaptic bouton fragmentation, accelerated neuronal aging, and locomotor decline. This toxicity is prevented in *Drosophila* by Turandot proteins, which reduce AMP–PS interactions on motoneurons. Together, our findings define a molecular mechanism underlying neuronal susceptibility to immunopathology and a set of proteins with neuroprotective potential.

**Graphical abstract:** This study shows that infection or dysbiosis in *Drosophila* can simultaneously induce antimicrobial peptide expression while promoting phosphatidylserine (PS) exposure on neurons at the neuromuscular junction. Cationic antimicrobial peptides contribute to neurodegeneration by binding to neurons that expose negatively charged phospholipids such as PS. In *Drosophila*, a family of secreted peptides, the Turandot proteins, can protect the peripheral nervous system by binding to PS-exposed membranes. Together, these findings reveal a role for antimicrobial peptides, a key component of innate immunity, in promoting neurodegeneration as well as a potential protective mechanism by PS masking agent.

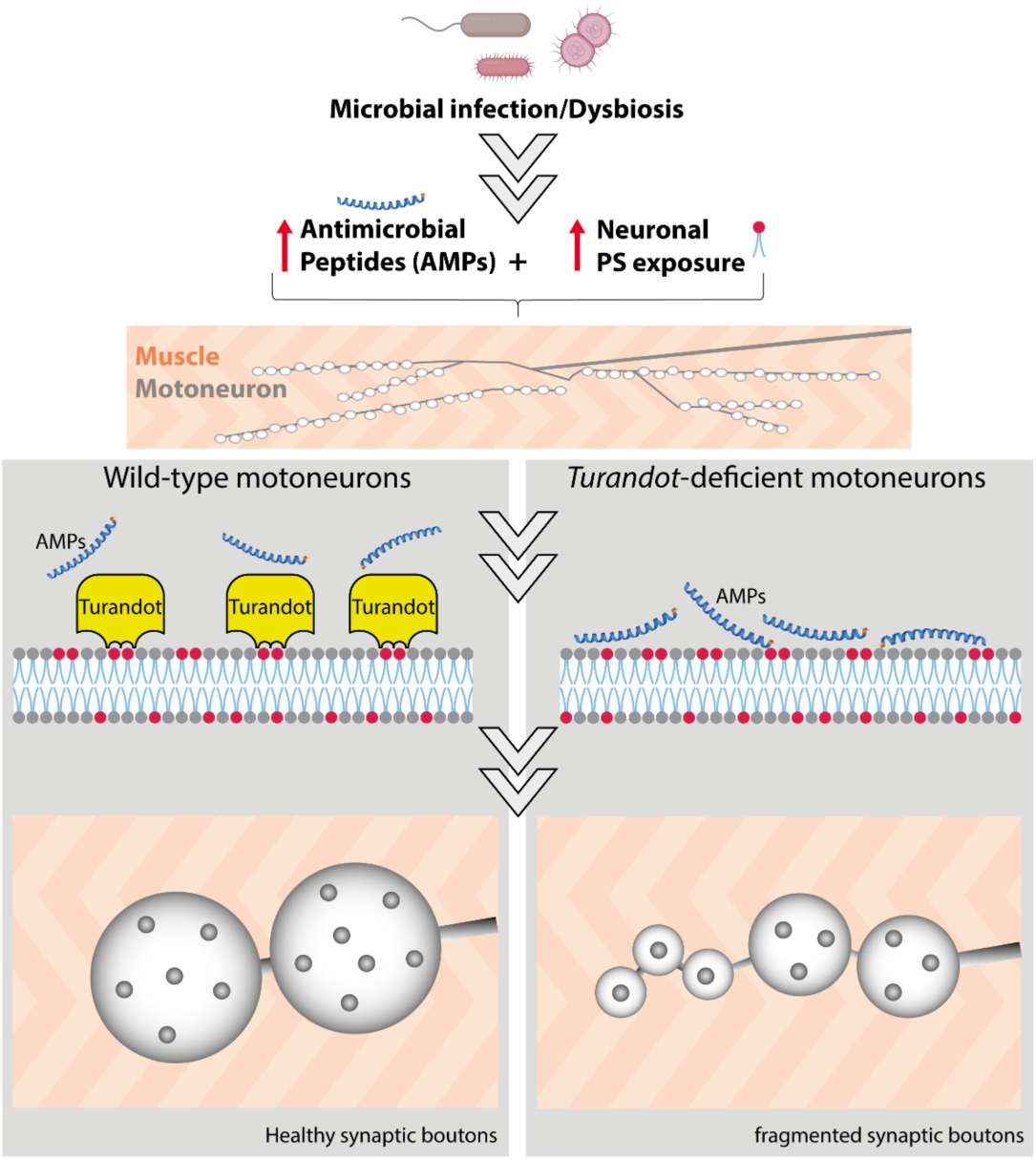

## Introduction

The immune and nervous systems are intimately interconnected. Once considered an immune-privileged site, the nervous system is now recognized as being actively shaped and maintained by immune mechanisms throughout life^1,2^. Immune cells and soluble effectors play critical roles in neural development and homeostasis, eliminating excess neurons, pruning synapses, and conveying information about infection and metabolic status to the brain^3–5^. At the same time, neuroinflammation, whether driven by infection or sterile insults, has emerged as a key contributor to brain aging and cognitive decline across a range of neurodegenerative disorders^6,7^. For example, Alzheimer’s disease is often associated with herpesvirus infections, and proteins implicated in pathogenic plaque formation in both Alzheimer’s and Parkinson’s diseases such as Amyloid-ß and α-synuclein have been reported to exhibit antimicrobial properties^8–11^.

Antimicrobial peptides (AMPs) are universal innate immune effectors that contribute to host defense in plants and animals^12,13^. They are usually small, cationic peptides with potent membranolytic and microbicidal activities against bacteria and fungi^14,15^. AMPs play a central role in host defense by preventing microbial infection at barrier epithelia and within host compartments. They also shape and regulate non-pathogenic microbes, including the gut microbiota^16,17^. In mammals, some AMPs further act as chemoattractants or immunomodulatory molecules, illustrating their broad functional diversity^18^. Owing to their positive charge, AMPs preferentially interact with microbial membranes, which typically have a strongly negative charge whereas host cell membranes are comparatively less negative. This difference in membrane net charge enables the immune system to selectively target infectious non-self while largely sparing host tissues^19,20^. However, recent evidence revealed that AMPs can interact with host tissues and exhibit cytotoxic activity, particularly toward tumor cells^21–25^.

In many organisms, AMP expression increases not only during infection but also with age^26–28^. Seminal studies in *Drosophila* demonstrated that AMP overexpression or chronic immune activation in the brain leads to neurodegeneration and shortened lifespan, revealing the intrinsic neurotoxicity of AMPs^29–32^. However, how AMPs influence the nervous system under physiological conditions, and the molecular mechanisms underlying AMP toxicity, remain largely unknown.

Eukaryotic plasma membranes are asymmetric in both lipid composition and biophysical properties^33–35^. Phosphatidylserine (PS), an abundant, negatively charged phospholipid, is typically restricted to the inner leaflet, making the outer leaflet electrically near-neutral^36,37^. Membrane phospholipid asymmetry is tightly regulated by the opposing activities of flippases, which translocate lipids from the outer to the inner leaflet of the membrane maintaining asymmetry, and scramblases, which redistribute them between both leaflets^38,39^. During apoptosis, scramblase activation collapses membrane asymmetry and drives PS externalization, marking cells for immune recognition and phagocytic clearance^5,40–43^. Transient and spatially confined PS exposure also occurs in healthy cells, where it regulates developmental and stress-response processes^44–47^. We and others recently showed that cells exposing PS can be targeted by cationic AMP, both in tumoral context and in healthy epithelia^22,23,48^. Because PS is highly enriched in the brain and is essential for neuronal survival and neurotransmission, PS exposure could represent a key determinant of AMP-mediated neurotoxicity.

The central nervous system is protected from circulating immune factors by the blood–brain barrier, whereas the peripheral nervous system is more directly exposed and thus potentially more susceptible to immune-mediated damage. The *Drosophila* neuromuscular junction (NMJ) provides a powerful model for investigating neurodevelopment, degeneration, and aging^49–52^. As in other organisms, locomotor performance declines with age in *Drosophila* and correlates with the fragmentation of synaptic boutons at the NMJ^53–55^. Bouton fragmentation thus serves as a sensitive and tractable readout of neuronal degeneration, offering a suitable platform to study AMP–neuron interactions.

We have previously shown that a family of stress-induced circulating proteins, the Turandots (Tots), can bind to PS and other negatively charged phospholipids to protect cells from AMPs^48^. Notably, Tots bind tracheal cells that constitutively expose high levels of PS. In *Tot*-deficient animals, cationic AMPs bind to exposed PS via electrostatic interactions and kill tracheal tissues, revealing a concealed toxicity of AMPs to host tissues that is ordinarily mitigated by Turandot resilience factors^48^. Here, we investigate how AMPs affect the peripheral nervous system during infection and aging. We show that AMPs bind to and damage motoneurons in a PS-dependent manner, and that this toxicity is normally buffered by Turandot proteins. In their absence, AMPs trigger NMJ degeneration during infection and drive premature neuronal terminal structural aging. Collectively, our study reveals a role of AMPs and PS exposure in neurodegenerative contexts.

## Results

### AMPs are detected at the neuromuscular junction

Several reports have proposed a role of AMPs in neurodegeneration^27,29,31^. We first asked whether immune-inducible AMPs physically interact with the peripheral nervous system. To this end, we generated a fly line carrying an endogenous knock-in of the antibacterial peptide gene *Diptericin*, tagged with HA (referred to as DptA KI). We then systemically infected these flies with the Gram-negative bacterium *Pectobacterium carotovorum carotovorum* 15 (*Ecc15*), a strong inducer of the Imd pathway that regulates AMP gene expression^56–58^. Following infection, we observed a marked accumulation of DptA-HA at the neuromuscular junction (Figure 1A). DptA staining on motoneurons could arise from neuronal expression, paracrine production by surrounding glia, or remote secretion from the fat body, the primary source of AMPs after systemic infection. Supporting the latter possibility, ectopic expression of an *UAS-DptA-HA* using the fat body driver Lpp-Gal4 (*Lpp>DptA-HA*) produced a motoneuron staining pattern comparable to that observed with *DptA KI* flies upon bacterial infection (Figure 1B). Similar results were obtained upon overexpression of two other AMP genes, *Drosomycin* and *Defensin*, indicating that several AMPs can bind motoneurons (Figure S1A). Conversely, reducing expression of DptA-HA KI by RNAi silencing the gene encoding the Imd pathway transcription factor Relish (Rel^56^) specifically in the fat body abolished DptA-HA binding to neurons upon bacterial infection (Figure 1C). Collectively, these experiments indicate that AMPs produced by the fat body upon infection can bind to motor neuron terminals.

**Figure 1:**
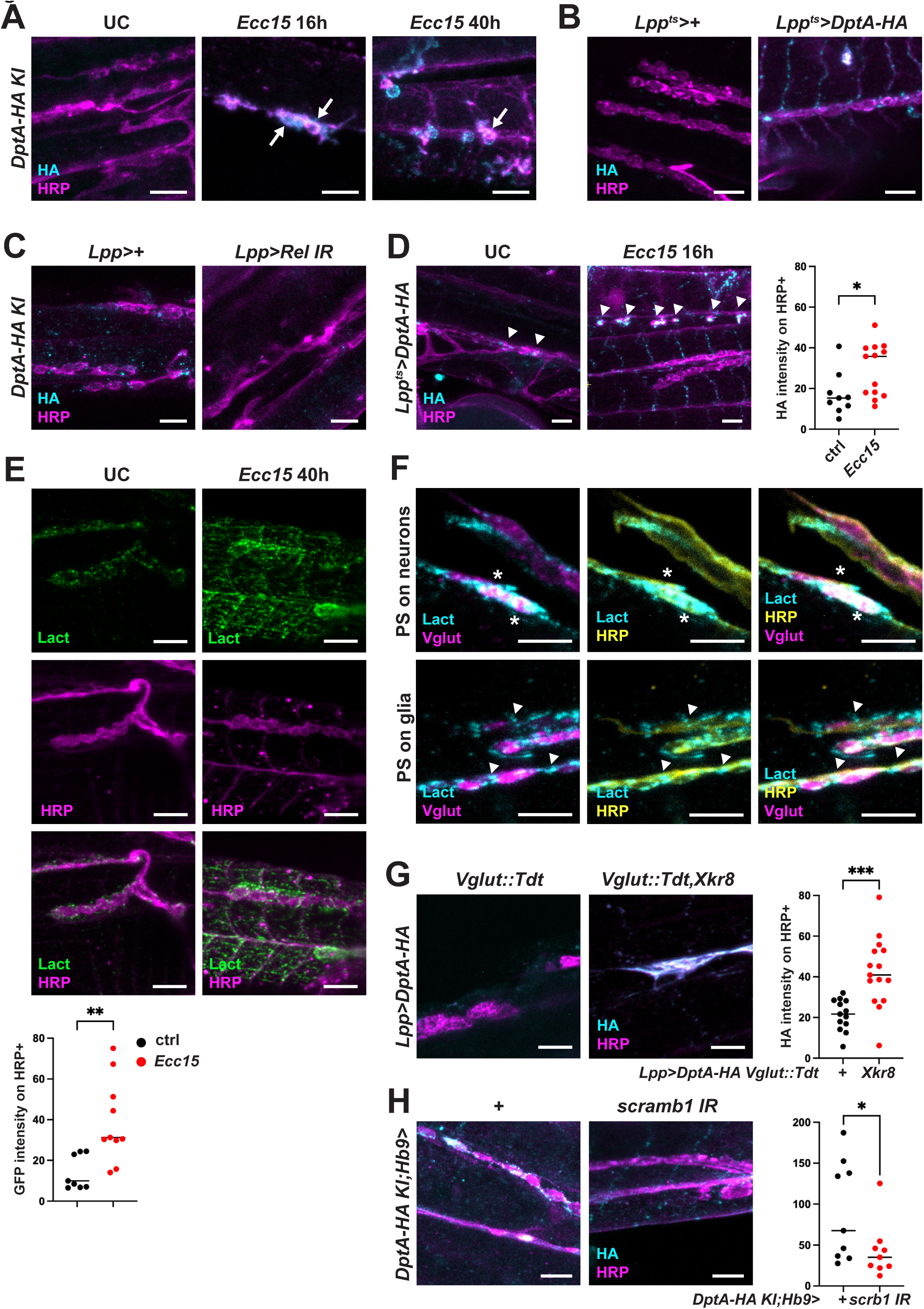
Antimicrobial peptides (AMPs) bind to exposed phosphatidylserine (PS) at the neuromuscular junction (NMJ) (**A-D**) Representative images of a NMJ stained for HA-tagged DptA-HA (cyan) and the presynaptic compartment (magenta, anti-HRP). (**A**) *DptA-HA KI* flies unchallenged (UC) or systemically infected with the gram-negative bacteria *Ecc15* for 16h or 40h at 29°C. White arrows indicate AMP colocalization with HRP. (**B**) NMJ staining of control (left) or flies overexpressing DptA-HA in the fat body (*Lpp^ts^>DptA-HA*). (**C**) NMJ staining of *DptA-HA KI* flies (left) or *DptA-HA KI* flies expressing an RNAi against *Relish* in the fat body (right) infected with heat-killed *Ecc15* for 16h at 29°C. (**D**) NMJ staining of flies overexpressing DptA-HA in the fat body (*Lpp^ts^>DptA-HA*) in unchallenged (UC, left) conditions or after 16h of *Ecc15* infection (right). Right panel: quantification of DptA-HA fluorescence intensity on HRP-positive structures in UC (black) and infected (red) conditions. (**E**) NMJ staining of flies expressing the PS externalization sensor GFP-lactadherin in the fat body (*Lpp^ts^>GFP-lact*) in unchallenged (left panels) and *Ecc15*-infected conditions (right). Green, GFP-lact; magenta, HRP. Histogram: quantification of GFP fluorescence intensity on HRP-positive structures in UC (black) and infected (red) conditions. (**F**) Representative images of *Ecc15*-infected flies expressing *Lpp^ts^>GFP-lact* (cyan) with TdTomato-labeled motoneurons (vglut-LexA::LexOp-TdTomato, magenta) and stained with anti-HRP (yellow). top panels show colocalization of PS with motoneurons (asterisks, HRP and TdTomato-positive). lower panels show co-staining of PS with glia (arrowheads, HRP-positive, TdTomato-negative). (**G**) Representative images of NMJ of *Lpp>DptA-HA* flies harboring TdTomato-labeled motoneurons (*vglut-LexA::LexOp-TdTomato*; left panel) or TdTomato-labeled motoneurons with increased PS exposure (*vglut-LexA::LexOp-TdTomato,xkr8*; right panel). Magenta, HRP; cyan, HA tag. Right histogram, quantification of DptA-HA fluorescence intensity on TdTomato-positive structures. (**H**) Representative images of NMJ of *DptA-HA KI* flies harboring control motoneuron driver (*Hb9>+*; left panel) or scramb1 RNAi driven in motoneurons (*Hb9>scramb1 IR*; right panel). Magenta, HRP; cyan, HA tag. Scale bars: 5 μm. Right histogram, quantification of DptA-HA fluorescence intensity on HRP-positive structures. Histograms: horizontal bar indicates the median and each dot represents an independent animal. Statistics: Mann-Whitney test; * for p between 0.01 and 0.05; ** for p between 0.001 and 0.01, *** for p < 0.0001.

### Secreted AMPs bind to PS exposing neuromuscular junctions

Interestingly, we observed that *lpp>DptA-HA* flies showed increased DptA recruitment to motoneurons following infection compared to unchallenged flies (Figure 1D). We have recently shown that AMPs bind cell membranes exposing phosphatidylserine (PS)^48^. This prompted us to examine whether there is an increased PS exposure on neurons upon systemic infection. To test this, we over-expressed the secreted PS binding sensor Lactadherin-GFP (Lact-GFP) using the *Lpp-Gal4* driver^59,60^. Extracellular PS staining with this sensor revealed elevated PS exposure at NMJ terminals after infection compared to the unchallenged condition (Figure 1E). Lact-GFP staining was observed in both the presynaptic (neuron and glia, HRP-positive) and postsynaptic compartment (muscle, HRP-negative) (Figure 1E). Co-staining of the presynaptic compartment with HRP and Vglut-LexA::LexO-TdTomato (*Vglut::Tdt*) revealed that both neurons (HRP and Tdt double-positive) and surrounding glia (HRP positive and Tdt negative) had Lact-GFP signals (Figure 1F). We therefore hypothesized that the increased exposure of PS at the NMJ upon systemic infection could explain the enhanced AMP binding observed in these animals. To test this directly, we increased PS exposure in motoneurons by overexpressing the Xkr8 scramblase (*Vglut::tdt,Xkr8*) in motoneurons of unchallenged flies expressing DptA-HA from the fat body (*Lpp>DptA-HA*). Elevating PS exposure in motoneurons of uninfected flies markedly increased the binding of DptA-HA to NMJ terminals (Figure 1G). Conversely, reducing PS exposure by knocking down the scramblase *scramb1* in a subset of motoneurons (*Hb9>scramb1 IR*) was sufficient to reduce DptA-HA recruitment to the NMJ terminals of these neurons (Figure 1H). Together, these findings indicate that systemic infection increases PS exposure on motoneurons allowing the binding of AMPs.

### Turandot proteins prevent infection-induced peripheral neuron fragmentation

Having established that AMPs can reach peripheral neurons upon systemic infection, we next asked whether this interaction causes motoneuron damage. To assess neuronal integrity, we quantified synaptic bouton fragmentation, a sensitive marker of neuronal injury and aging^61^. In young adults, motoneuron terminals are composed of a chain of synaptic boutons each of which contain multiple active zones that mediate neurotransmitter release. In tandem with aging, the number of active zones per bouton as well as the average diameter of bouton decrease^53,54^. This degenerative process results in the accumulation with age of synaptic boutons with only a single active zone, termed fragmented boutons^53,54^. We therefore quantified the proportion of motoneuron boutons (labeled with the OK6-gal4 driver) that were fragmented using the active zone marker Bruchpilot (Brp). To exclude effects from bacterial virulence, we used a cocktail of heat-killed (HK) *P. carotovorum 15 and Micrococcus luteus* bacteria, allowing us to specifically test the impact of the immune response. Injection of dead bacteria in wild-type flies did not induce any major changes in synaptic morphology causing only a slight increase in single-active zone boutons four days post-infection. This indicates that AMPs are not overtly neurotoxic in infected wild-type flies (Figure 2A and 2B). We previously showed that the stress-induced Turandot (Tot) proteins protect PS-exposing tracheal cells from AMP toxicity^48^. The *Drosophila* genome encodes eight Tots that are secreted in the hemolymph upon infection^62,63^. Turandots bind to PS on the cell surface preventing AMP-induced pore formation. We therefore hypothesized that Turandot proteins could similarly shield neurons from AMP toxicity during infection. Use of anti-TotA antibody revealed TotA enrichment at the NMJ following infection (Figure 2C). We thus tested if *Tot*-deficient neurons were more affected upon immune stimulation. While neurons of wild-type animals remained unaltered, infected *Tot*-deficient flies showed a strong increase in the proportion of fragmented single active zone boutons (Figure 2A, white arrowheads and 2B). This phenotype depended on immune activation, as unchallenged wild-type and *Tot*-deficient animals did not exhibit additional fragmented boutons (Figure 2A and 2B). Similar results were obtained in flies lacking either five (*Tot^AZX^* flies, lacking TotA, B, C, X and Z,) or six (Tot*^XMAZ^* flies, lacking TotA, B, C, M, X and Z) out of the eight *Turandot* genes (Figure 2A and 2F). Having shown that Tots prevent motoneuron terminal degeneration upon infection, we then tested if *Tot* deficiency affected AMP binding to neurons. Infected *Tot* deficient flies showed a stronger recruitment of DptA and CecA to the NMJ (Figure 2D and 2E), consistent with the notion that Tots prevent AMP binding to PS-exposing neurons. To determine whether AMPs could induce terminal degeneration, we generated flies simultaneously lacking Tots and 12 AMPs, including two Diptericins, three Attacins, four Cecropins, one Drosocin, one Defensin, and one Metchnikovin (Δ*AMP12, Tot^AZX^*)^14,48,64^. Removal of AMPs in *Tot^AZX^* flies completely rescued the number of active zones per bouton upon infection, demonstrating that AMPs act to enhance motoneuron terminal fragmentation in this background (Figure 2F). Deleting only the four *cecropin* genes (*CecA1/A2, B* and *C*) yielded a partial rescue, indicating that Cecropins substantially contribute to motoneuron terminal fragementation in absence of *Tot* (Figure 2F). Together, our study reveals that AMPs induced upon systemic infection can damage the peripheral nervous system, and that Turandot proteins can protect motoneurons from this toxicity by preventing AMP binding to NMJ terminals.

**Figure 2:**
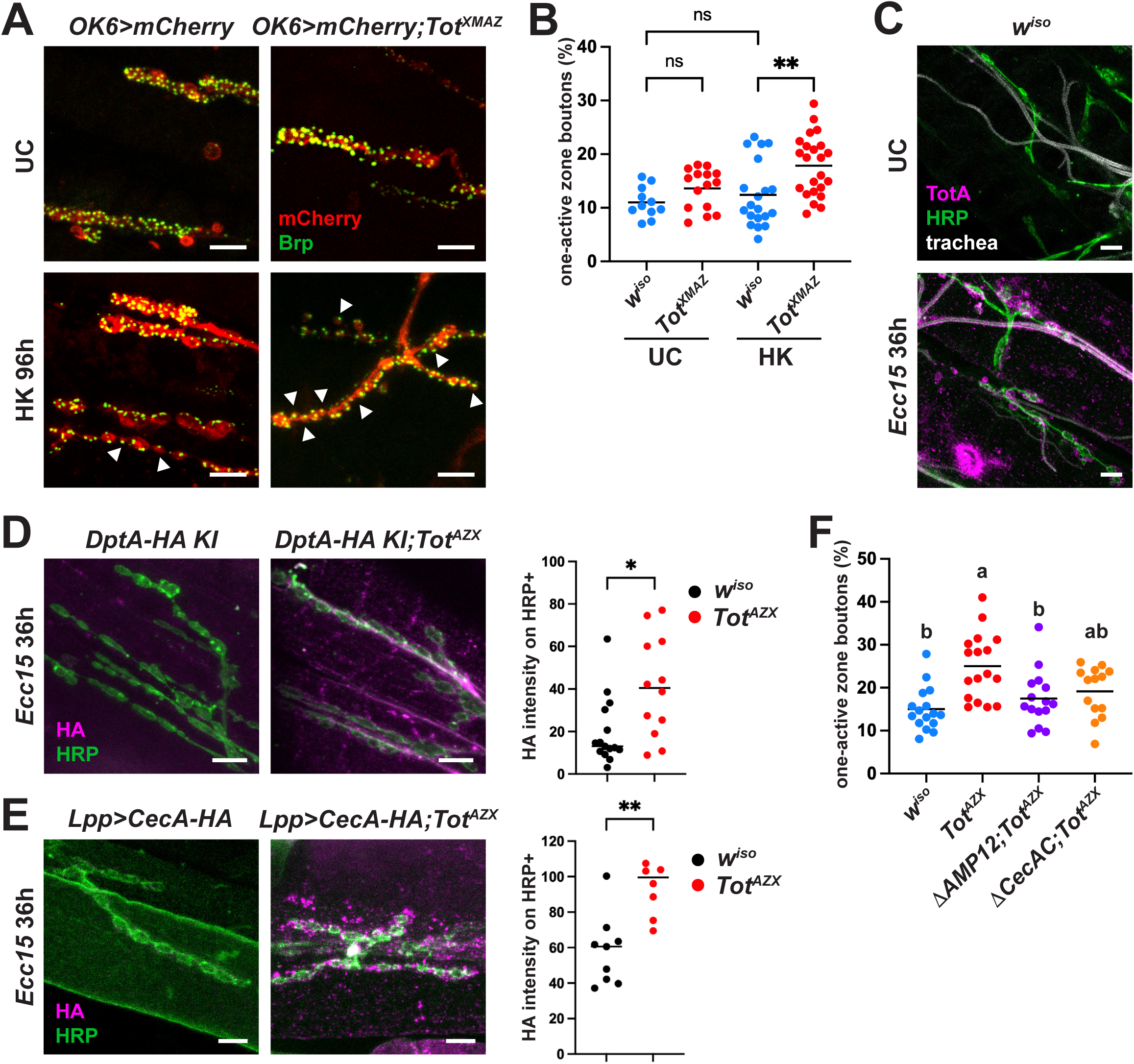
Turandot proteins prevent infection-induced synaptic bouton fragmentation. (**A**) NMJ of control (*OK6>mCherry*, left) or Tot deficient (*OK6>mCherry;Tot^XMAZ^*, right) flies unchallenged (UC, top) or infected with heat-killed bacteria (*Ecc15* HK, bottom). Motoneurons are stained in red and the active zone marker Brp in green. Arrowheads indicate fragmented boutons containing a single active zone. (**B**) Quantification of single-active zone synaptic boutons in *w^iso^* (blue) and *Tot^XMAZ^* (red) in control flies (UC) flies challenged with a mixture of heat-killed *Ecc15* and *Micrococcus luteus (HK)*. (**C**) NMJ staining of UC (top) or *Ecc15*-infected flies (bottom); white, chitin; green, HRP and magenta, TotA. (**D**-**E**) NMJ staining of *Ecc15*-infected control (left) or *Tot^AZX^*-deficient flies (right) carrying *DptA-HA KI* (D) or overexpressing CecA-HA in the fat body (E). Cyan, HA; Magenta, HRP. Right panels, quantification of HA fluorescence intensity in HRP-positive structures. Black, *w^iso^*; red, *Tot^AZX^*. (**F**) Quantification of single-active zone synaptic boutons in *w^iso^* (blue), *Tot^AZX^* (red), Δ*AMP12;Tot^AZX^* (purple) and Δ*CecAC;Tot^AZX^* (black) in heat-killed Ecc15-infected flies. Scale bars: 5 μm. Horizontal bars in histograms indicate the median and each dot represents an independent animal. Statistics: (D-E) Mann-Whitney test; (B,F) One-way ANOVA followed by a Dunnett’s multiple comparisons test; ns, not significant, * for p between 0.01 and 0.05; ** for p between 0.001 and 0.01, *** for p < 0.0001.

### Turandots proteins prevent motoneuron fragmentation upon aging

Aging is accompanied by a decline in motoneuron function in parallel with increased synaptic bouton fragmentation^54^. Having shown that motoneurons in *Tot*-deficient flies had enhanced fragmentation upon immune stimulation, we next asked whether these mutants exhibit precocious bouton fragmentation during aging. As noted above, neither young wild-type nor young *Tot*-deficient animals displayed motor neuron terminal defects. Strikingly, 20-day-old *Tot*-deficient flies already exhibited pronounced bouton fragmentation, whereas age-matched wild-type flies did not. By the age of 36 days, both wild-type and *Tot* mutants showed comparable levels of degeneration. This suggested an acceleration of aging during middle-age of motor neurons in *Tot*-deficient flies (Figure 3A and B). This phenotype was reproduced in another genetic background expressing mCherry in motoneurons (*OK6>mCherry*) and was comparable in 20-day-old *Tot^AZX^* and *Tot^XMAZ^* mutants (Figure 3C). Consistent with the idea that *Tot* mutants undergo premature NMJ terminal aging, overexpression of the guanine nucleotide exchange factor Trio, that preserves motoneuron architecture and delays aging, was sufficient to rescue bouton fragmentation in *Tot*^AZX^ mutants (Figure 3D)^53^. The accelerated age-related degeneration of *Tot* mutants correlated with reduced locomotor performance in a climbing assay (Figure 3E). This decline did not reflect an overall health deterioration, as lifespan was not altered in *Tot*-deficient flies (Figure 3F).

**Figure 3:**
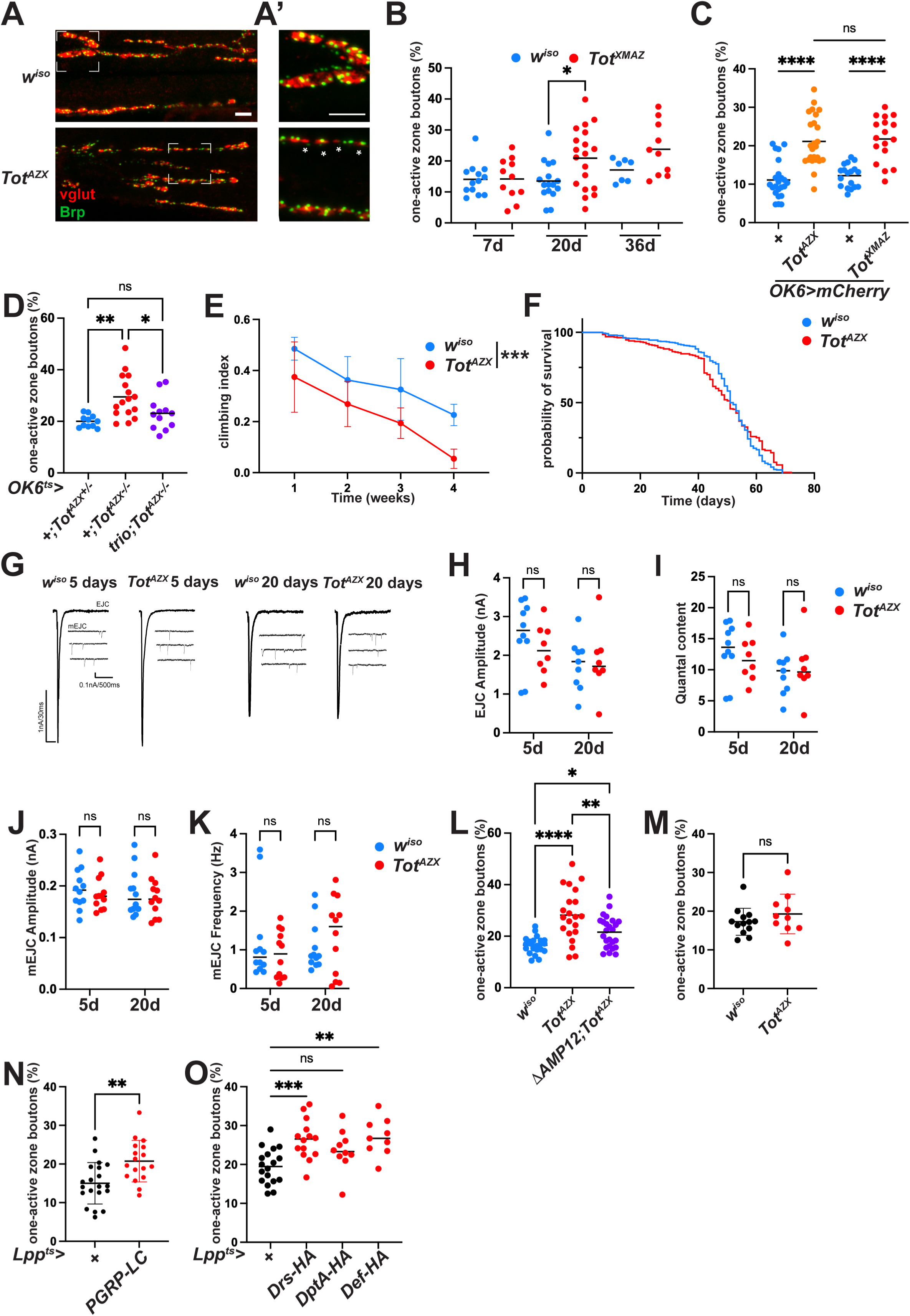
Turandot proteins protect against age-dependent motoneuron fragmentation. (**A**) Representative confocal micrographs of NMJ of 20 days old *w^iso^* (top) and *Tot^AZX^* flies (bottom) stained for Vglut (green) and Brp (red) (A’) Higher magnification of the area defined by the dotted square in (A). Scale bars: 5 μm. (**B**) Quantification of single-active zone bouton proportion in *w^iso^* (blue) and *Tot^XMAZ^* (red) at 7, 20 and 36 days. (**C**) Quantification of single-active zone bouton proportion in control (blue), *Tot^AZX^* (orange) and *Tot^XMAZ^* (red) flies with mCherry-labeled motoneurons. (**D**) Quantification of single-active zone bouton proportion in control (blue), *Tot^AZX^* (red) and *Tot^AZX^* overexpressing Trio in motoneurons (purple). (**E,F**) Climbing ability (E) and survival (F) as a function of time of *w^iso^* (blue), *Tot^AZX^* (red) flies. (**G**) Representative traces of excitatory junctional currents (EJC)(above) and miniature excitatory junctional currents (mEJC)(below) from 5 days old and 20 days old terminals. (**H-I**) Quantification of EJC amplitude (**H**) and quantal release (**I**) of terminals from 5 day- and 20 day-old *w^iso^* (blue) and *Tot^AZX^* (blue). (**J-K**) Quantification of mEJC amplitude (**J**) and frequency (**K**) of terminals from 5 day- and 20 day-old *w^iso^* (blue) and *Tot^AZX^* (blue). (**L-O**) Quantification of bouton fragmentation in control (blue), *Tot^AZX^* (red) and Δ*AMP12;Tot^AZX^* (L, purple) raised in conventional (**L**) or axenic medium (**M**). (**N,O**) Quantification of single-active zone boutons in 20 days old control flies (black) or flies overexpressing the Imd pathway receptor PGRP-LC (**N**, red) or the AMPs Drs, DptA or Def (**O**, red). Horizontal bar indicates the median and each dot represents an independent NMJ. Statistics: (B,M,N), Mann-Whitney test; (C,D,H-K,L,O), One-way ANOVA followed by a Dunnett’s multiple comparisons test; (E), Mixed-effects model with Geisser-Greenhouse correction; (F) Logrank (Mantel-Cox). ns, not significant for p > 0.05, * for p between 0.01 and 0.05; ** for p between 0.001 and 0.01, *** for p < 0.0001.

Age-dependent synaptic bouton fragmentation occurs in tandem with reduced neurotransmission^54^. Two forms of vesicular neurotransmission occur at the *Drosophila* neuromuscular junction (NMJ): evoked neurotransmission, in which multiple synaptic vesicles are released in response to an action potential, and miniature neurotransmission, which involves the release of single vesicles independent of action potentials^65,66^. Both forms decline with age, with miniature neurotransmission playing a critical instructive role in maintaining synaptic structural integrity^54^. Having established premature synaptic fragmentation in *Tot*-deficient flies, we next examined the functional properties of these synapses in both young and 20-day-old animals. In 20-day-old control flies we observed a decline of both forms of neurotransmission (Figure 3G-K)^54^. Evoked junctional current (EJC) amplitudes, miniature EJC (mEJC) amplitude, quantal content and mEJC frequency in *Tot* mutants was not different to these age-matched controls (Figure 3G-K). These data indicate that Tot mutants do not exhibit overt neurotransmission defects despite their altered synaptic architecture (Figure 3A-C), similar to Trio mutants^54^. The preservation of miniature neurotransmission in *Tot*-deficient flies suggests that bouton fragmentation in these mutants is unlikely to result from impairments in this process. Rather, these findings show that Turandots delay age-dependent bouton fragmentation via an independent mechanism.

### Turandots antagonize AMP toxicity to motoneurons

Based on our findings in the context of systemic infection, we hypothesized that AMPs contribute to synaptic bouton fragmentation in *Tot* mutants upon aging. Supporting this idea, deletion of 12 AMPs in *Tot*-deficient background partially restored bouton structures to wild-type levels in 20-day-old flies (Figure 3L). This result shows that AMPs can damage the peripheral nervous system not only during infection but also in the course of aging, prompting us to investigate their source in this context.

AMP expression increases with aging and this increase is largely driven by the microbiota, as flies raised in axenic conditions maintain lower levels of AMPs^26,27^. We therefore tested whether the microbiota contributes to the premature NMJ degeneration observed in *Tot* mutant flies by promoting increased AMP expression. Indeed, axenic *Tot-*deficient animals did not display the bouton fragmentation observed in 20-day-old conventionally raised *Tot*-deficient flies, demonstrating that the microbiota drives age-associated bouton decline in these mutants (Figure 3M). Together, these findings indicate that the premature aging observed in *Tot* mutants results from increased AMP expression induced by the microbiota. These results show that increased AMP expression during aging is required to induce bouton fragmentation in the absence of Tot proteins, and suggest that maintaining a balance between AMPs and Tots is critical to preserve synaptic integrity. To test this directly, we asked whether overexpressing AMPs in wild-type flies is sufficient to induce bouton fragmentation. Chronic immune activation by overexpression of *PGRP-LC* in the fat body (*Lpp>PGRP-LC*) was sufficient to induce precocious bouton fragmentation (Figure 3N). Similarly, the constitutive overexpression of a single AMP, Diptericin, Drosomycin or Defensin from the fat body was also sufficient to induce motoneuron terminal fragmentation in 20-day-old flies (Figure 3O), demonstrating that AMPs are intrinsically neurotoxic when produced in excess. Collectively, our study reveals that AMPs produced in response to microbial dysbiosis during ageing possess a neurotoxic potential that is suppressed by Turandot proteins.

### Age-related PS exposure drives AMP neurotoxicity

Having shown that AMPs accelerate motoneuron bouton fragmentation during aging, we next assessed the role of PS exposure in this process. Imaging NMJ terminals at different ages revealed a slight increase in PS exposure as flies aged, suggesting a role of PS in AMP sensitivity (Figure 4A). To test this hypothesis, we reduced PS exposure in *Tot*-deficient flies either by overexpressing the flippase ATP8A or by knocking down the *scramblase 1* gene. Because exposed PS was detected in both the presynaptic and postsynaptic NMJ terminal compartments, we lowered PS translocation in either muscles or motoneurons using specific Gal4 drivers. Reducing PS exposure in *Tot*-deficient flies specifically in motoneurons by both approaches was sufficient to suppress bouton fragmentation demonstrating that PS exposure is necessary for this phenotype (Figure 4B and 4C). In contrast, lowering PS exposure in the muscle compartment did not rescue fragmentation in *Tot* mutants, highlighting the specific contribution of neuronal PS exposure in this process (Figure 4D). A potential role for glia in bouton fragmentation has been suggested previously^54^. Moreover, exposed PS on neurons usually acts as an ‘eat-me’ signal recognized by the PS receptor Draper that triggers glial phagocytosis^67,68^. We thus tested whether inhibiting glial phagocytosis prevented bouton fragmentation in *Tot* mutant flies. Glial knockdown of *Draper* in *Tot*-deficient flies failed to rescue the increase level of bouton fragmentation, arguing against a role of glial phagocytosis in this context (Figure 4E).

**Figure 4:**
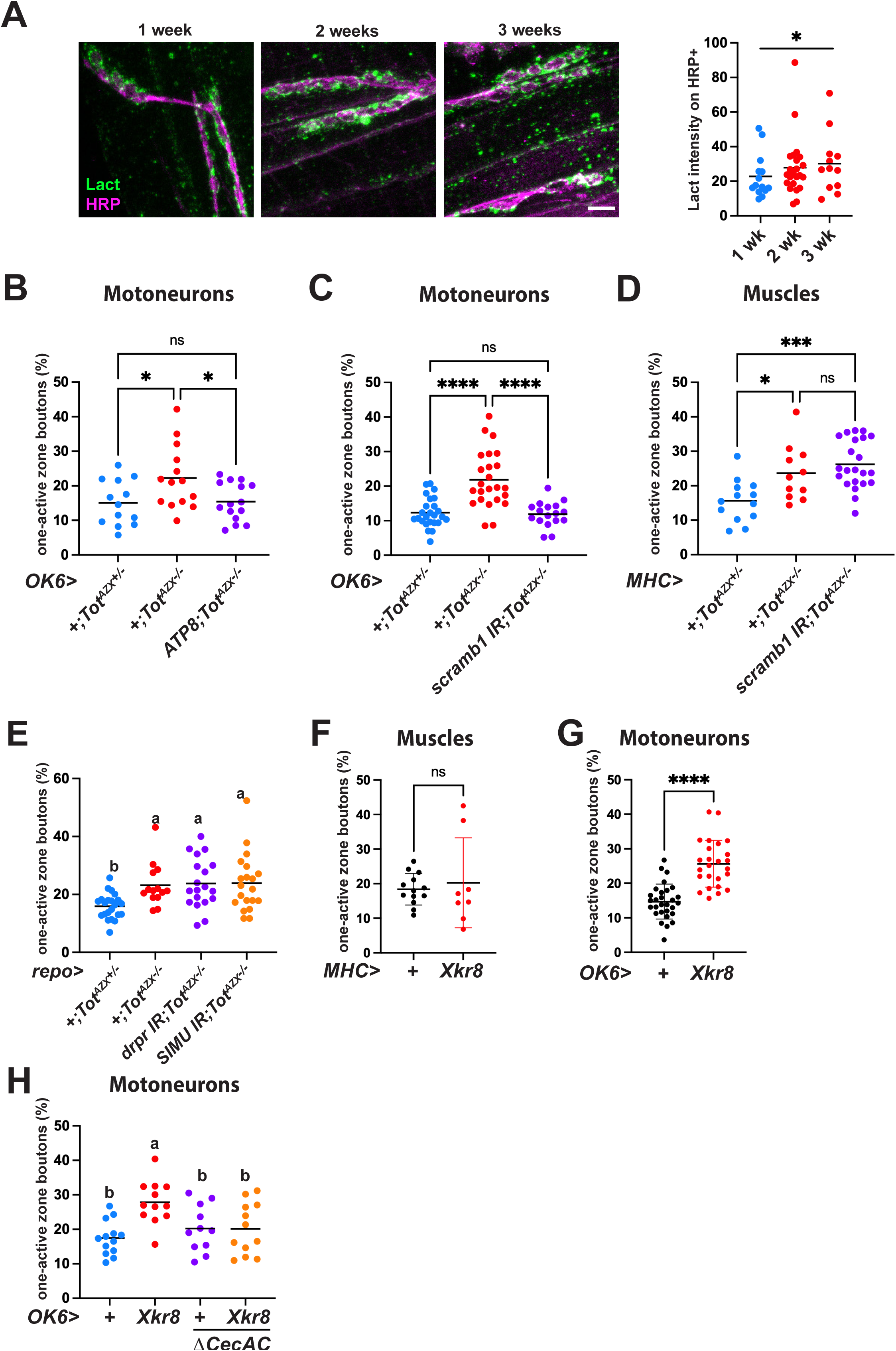
Age-related PS exposure drives AMP neurotoxicity. (**A**) Representative confocal micrographs of NMJ of 1-, 2- or 3-weeks old flies expressing GFP-lact in the fat body. Green, GFP; magenta, HRP. Scale bar: 5 μm. Right panel, quantification of GFP intensity on HRP-positive structures. (**B-D**) quantification of bouton fragmentation in 20 days old control (blue), *Tot^AZX^* (red) and *Tot^AZX^* animals with motoneurons expressing the ATP8 flippase (B, purple) or *Tot^AZX^* flies carrying an RNAi against *scramb1* expressed in motoneurons (C, purple) or muscles (D, purple). (**E**) Quantification of bouton fragmentation in control (blue), *Tot^AZX^* (red) and *Tot^AZX^* driving a *Draper* RNAi (purple) or a *SIMU* RNAi (blue) in glia. (**F-H**) Quantification of single-active zone bouton proportion in control animals (black) or flies expressing the Xkr8 scramblase in muscles (F, red) or in motoneurons (G, H, red) in a control genetic background (red) or in a Δ*CecAC* background (orange). Horizontal bar indicates the median and each dot represents an independent NMJ. Statistics: (F,G) Mann-Whitney test; (A-E,H) One-way ANOVA followed by a Dunnett’s multiple comparisons test; ns, not significant for p > 0.05, * for p between 0.01 and 0.05; ** for p between 0.001 and 0.01, *** for p < 0.0001; **** for p < 0.00001.

We then asked whether increasing PS exposure was sufficient to drive age-related fragmentation. Overexpression of the scramblase Xkr8 in neurons but not muscles significantly increased the proportion of fragmented boutons, revealing that neuronal PS exposure is sufficient to induce bouton fragmentation (Figure 4F and 4G). Strikingly, this effect was strictly dependent on Cecropins, as deletion of the *cecropin* locus fully suppressed PS-driven fragmentation in flies overexpressing *Xkr8* (Figure 4H). Together, these findings show that age-associated PS exposure in motoneurons licenses AMP binding and toxicity, thereby promoting the structural degeneration of NMJ terminals during aging.

## Discussion

Inflammation is a hallmark of aging and is implicated in age-associated neurological disorders^6,7,69^. Soluble immune factors such as reactive-oxygen species and complement components can damage neurons, either directly through membranolytic activity, or indirectly by impairing mitochondria or activating bystander glia^5,70,71^. Here, we describe antimicrobial peptides (AMPs) as a previously underappreciated neurotoxic effectors of inflammation. We show that AMPs produced in response to bacterial exposure can bind to and damage neurons of the peripheral nervous system. This phenomenon is limited in wild-type flies by Turandot proteins that protect host tissues by masking negatively charged phospholipids. Consistent with this, *Turandot* deficiency exacerbates AMP neurotoxicity in the context of immune response and aging. Our data show that AMP binding depends on phospholipid scrambling in the motoneuron plasma membrane. We hypothesize that cationic peptides can interact electrostatically with negatively charged phosphatidylserine (PS) exposed at the plasma membrane of neurons. While AMPs have previously been implicated in neurodegeneration indirectly, our study provides an in vivo demonstration that AMPs can directly bind to and damage neurons. We propose that PS exposure in non-apoptotic contexts and cationic antimicrobial peptides contribute to neurodegenerative diseases.

Exposed PS is traditionally described as an ‘eat-me’ signal that triggers engulfment by phagocytic glia during nervous system development. Instead, our study reveals that AMPs directly contribute to synaptic terminal fragmentation. Consistent with this, Draper is not involved in the bouton fragmentation phenotype of *Tot*-deficient adult flies. How AMP binding to motoneuron leads to bouton fragmentation remains unclear. Reduced miniature neurotransmission precedes synapse fragmentation and some AMPs have been shown to regulate sleep and memory, possibly through an effect on neurotransmission^72,73^. It would be tempting to speculate that AMP lowers miniature transmission, leading to bouton fragmentation. However, our data do not support this model, since we could not detect any reduction in miniature neurotransmission frequency in *Tot*-deficient animals. The fact that Trio overexpression preserves synaptic integrity in *Tot* mutant flies favors a model in which AMPs may directly damage neurons. The amphipathic properties of these peptides may destabilize the neuronal lipid bilayer through pore formation or detergent-like activity, leading to synaptic terminal damage. Alternatively, we cannot exclude that AMPs may also translocate across the membrane without overt disruption and act on intracellular targets, such as ribosomes or mitochondria^74,75^. Elucidating how AMP binding to the plasma membrane leads to synaptic terminal fragmentation will require further mechanistic investigation.

Immune induction alone does not seem sufficient to trigger NMJ terminal bouton fragmentation as this requires both AMP expression and neuronal PS exposure. Our data is in line with a two-hit model in which only vulnerable neurons, i.e. neuron exposing PS, are susceptible to AMPs triggered in response to a microbial infection. AMPs are induced upon infection through the Imd-NF-κB immune pathway, that has already been implicated in multiple neurodegenerative contexts^6,27,29,31,69^. During aging, gut bacterial overgrowth and dysbiosis promote chronic and sustained AMP expression by the gut and the fat body^26,76–78^. These small soluble peptides released in the hemolymph could diffuse through the synaptic cleft, bind to membranes of neurons exposing PS, leading to terminal degeneration. Consistent with this model, removing AMPs or reducing their expression by antibiotic treatment is sufficient to fully rescue the synaptic degeneration observed in 20 days old *Turandot-*deficient flies. Future work should identify the microbial species driving this response and determine whether their effects are mediated solely through AMP upregulation. Given the established links between gut microbiota alterations and neurodegenerative diseases such as Alzheimer’s, Parkinson’s, and Huntington’s diseases, assessing AMP contributions in these contexts will be of particular interest^79^.

The mechanisms driving neuronal PS exposure during infection and aging remain poorly understood. Neurons are particularly enriched in PS, which plays essential roles in metabolism, membrane dynamics, and cell survival^80^. The level of PS exposure is tightly regulated by a complex set of flippases and scramblases^39,81^. Under conditions of infection or aging, cellular stress or senescence may disrupt the balance of scramblase and flippase activities, leading to a collapse of membrane lipid asymmetry and consequent PS externalization. The same enzymes regulating phospholipid asymmetry are also involved in synaptic homeostasis, where they regulate receptor localization, vesicle fusion rates, and membrane dynamics^82–84^. Thus, PS exposure may not only serve as a marker of cellular stress, but also reflect active membrane remodeling processes intrinsic to neuronal plasticity. Determining whether PS exposure arises from a failure to maintain lipid asymmetry under stress or represents an adaptive response to such stress will be an important area for future investigation.

Finally, we demonstrate that Turandot proteins protect neurons from AMP-induced neuronal damage. Building on prior in vitro findings that Turandots bind PS and inhibit AMP-mediated pore formation, our study reveals a protective role for Turandots in the *Drosophila* peripheral nervous system^48^. In wild-type flies, AMPs are not overtly toxic to motoneurons during infection and AMP deficiency does not improve lifespan or motor performance^85^ AMP neurotoxicity becomes evident only in the absence of Turandot proteins or upon chronic AMP overexpression, indicating that a balanced expression of Turandots and AMPs is required to maximize antimicrobial defense while limiting collateral damage. In line with this, and consistent with a competition between AMPs and Turandots for PS binding, the loss of Turandots enhances neuronal binding of DptA and CecA. Without Turandot protection, AMP-PS interactions lead to precocious terminal structural fragmentation and locomotor defects. Whether similar protection operates in other neuronal subtypes, such as sensory neurons, remains to be determined. The fact that *Drosophila* evolved eight Turandot genes that are expressed in multiple tissues along its life cycle underscores AMP intrinsic toxicity to host tissues. Although Turandot genes are specific to *Drosophila*, both AMPs and PS translocation mechanisms are evolutionarily conserved, raising the possibility that analogous protective factors exist in other species, safeguarding the nervous system from AMP-induced membrane damage.

## RESOURCE AVAILABILITY

### Lead contact

Bruno Lemaitre

### Materials availability

Reagents generated in this study are available upon request to the academic community. For further inquiries, please address to the lead contact.

## ACKNOWLEDGMENTS

We thank the Bloomington Stock Center (USA), the Vienna Drosophila Resource Center and FlyORF (Switzerland) for fly stocks; Chun Han for providing fly strains; the BioImaging & Optics Platform (BIOP) in EPFL for confocal microscopy. We are very grateful to Florent Masson for comments and editing on the manuscript. This work was supported by the Swiss National Science Foundation grants SNSF TMAG-3_225935 to B.L. and 31003A_179587 and 320030-232324 to B.M.

## AUTHOR CONTRIBUTIONS

SR and BL conceived and designed the study. SR performed and analyzed most experiments. SW performed and analyzed the climbing assays. JPB generated the transgenic fly lines and performed the immunostainings. SV performed and analyzed the electrophysiology experiments. KK performed bacterial injections. BDM provided key resources. BDM and BL acquired funding. BL and SR administered the study. BL supervised the study. SR and BL wrote the initial draft and SV and BDM reviewed and edited the manuscript.

## DECLARATION OF GENERATIVE AI AND AI-ASSISTED TECHNOLOGIES in the writing process

During the preparation of this work, the authors used ChatGPT in order to correct English grammar. After using this tool or service, the authors reviewed and edited the content as needed and take full responsibility for the content of the publication.

## Supplementary Figure legends

**Figure S1:**

(A) Representative confocal images of a NMJ of control flies (left), or flies overexpressing Defensin (Def, middle panel) or Drosomycin (Drs, right panel) stained for HA-tagged DptA-HA (cyan) and the presynaptic compartment (magenta, HRP). Scale bars: 5 μm.

**Figure S2:**

(A,B) Quantification of input resistance (A) and resting membrane potential (B) of 5 day- and 20 day-old A2 mvim terminals from *w^iso^* (blue) and *Tot^AZX^* (blue).blue

## Star Methods

### Key resources table

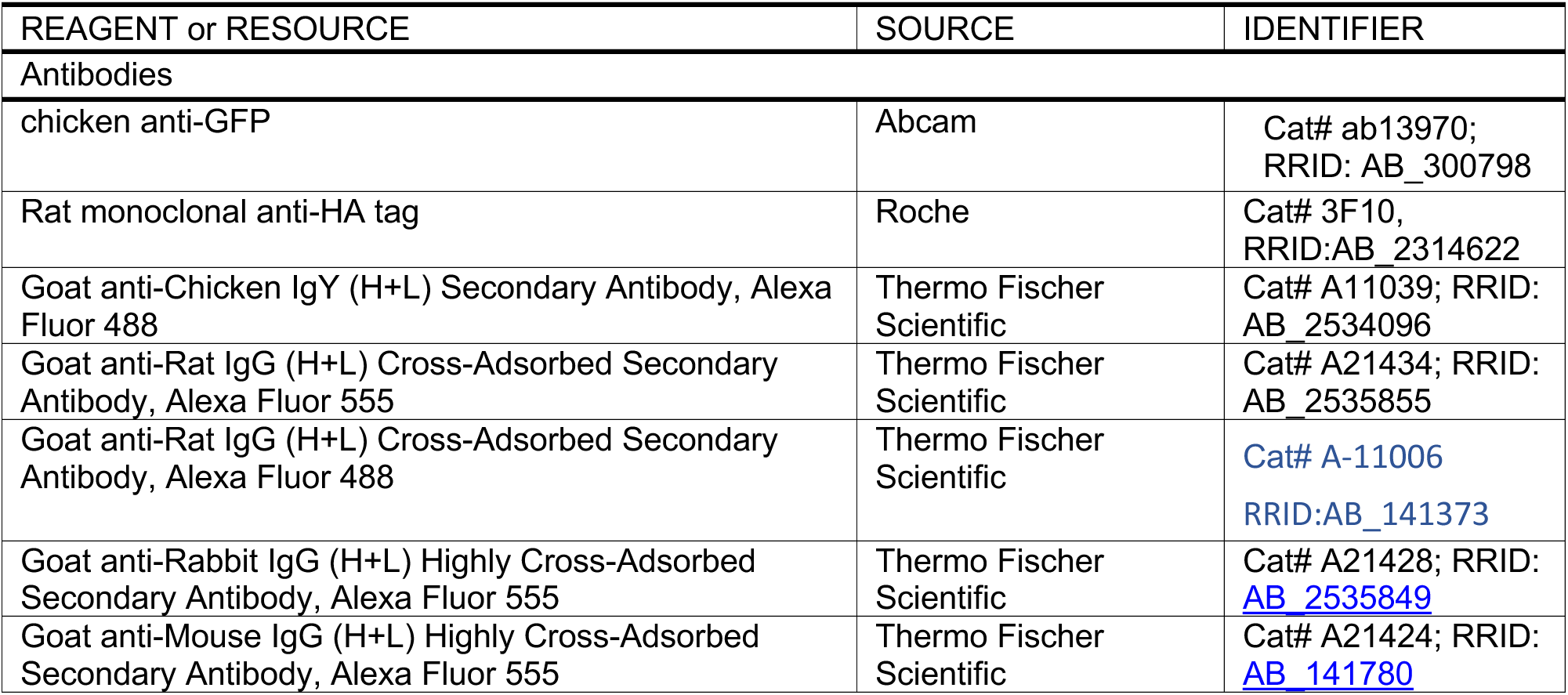
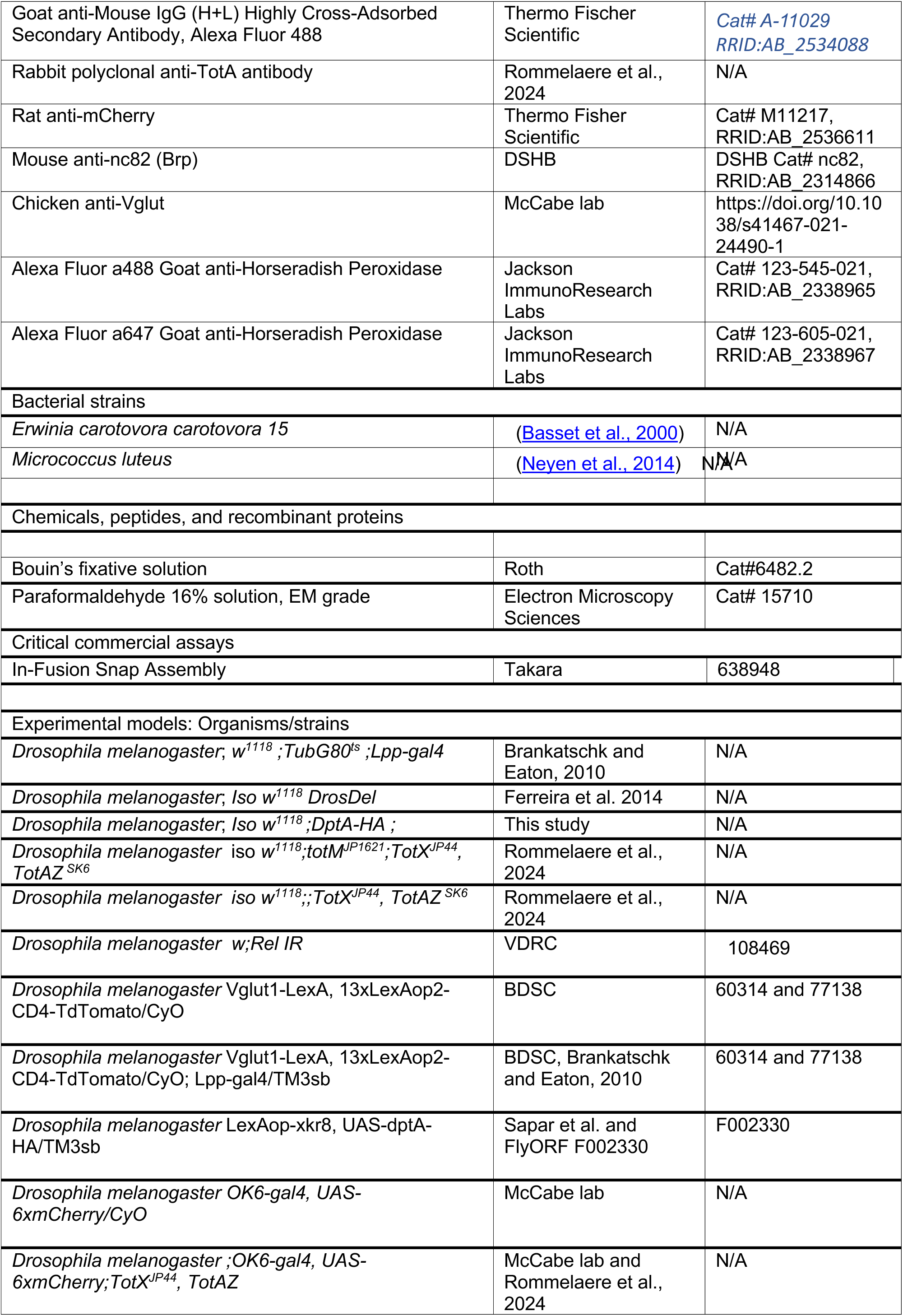
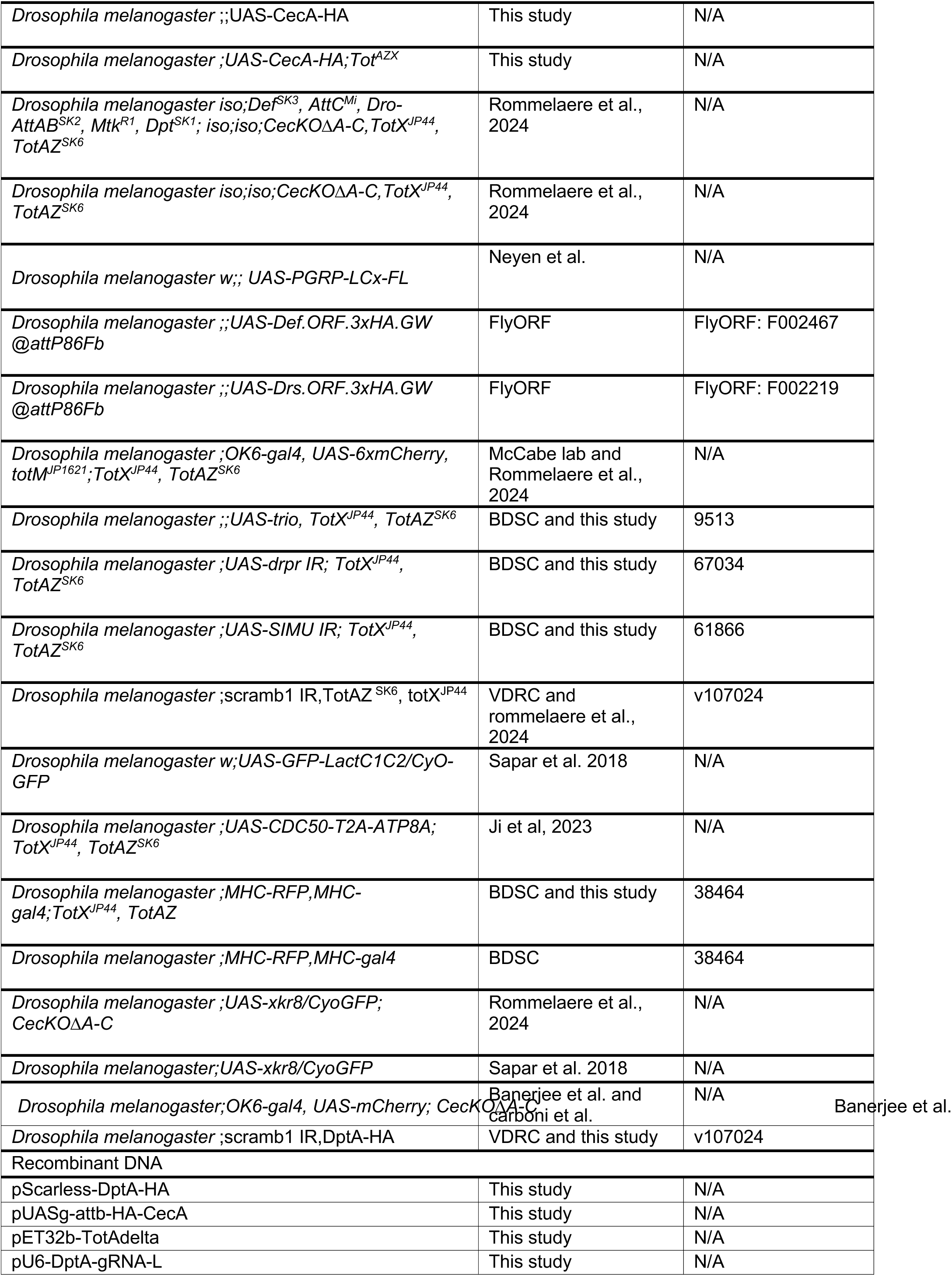
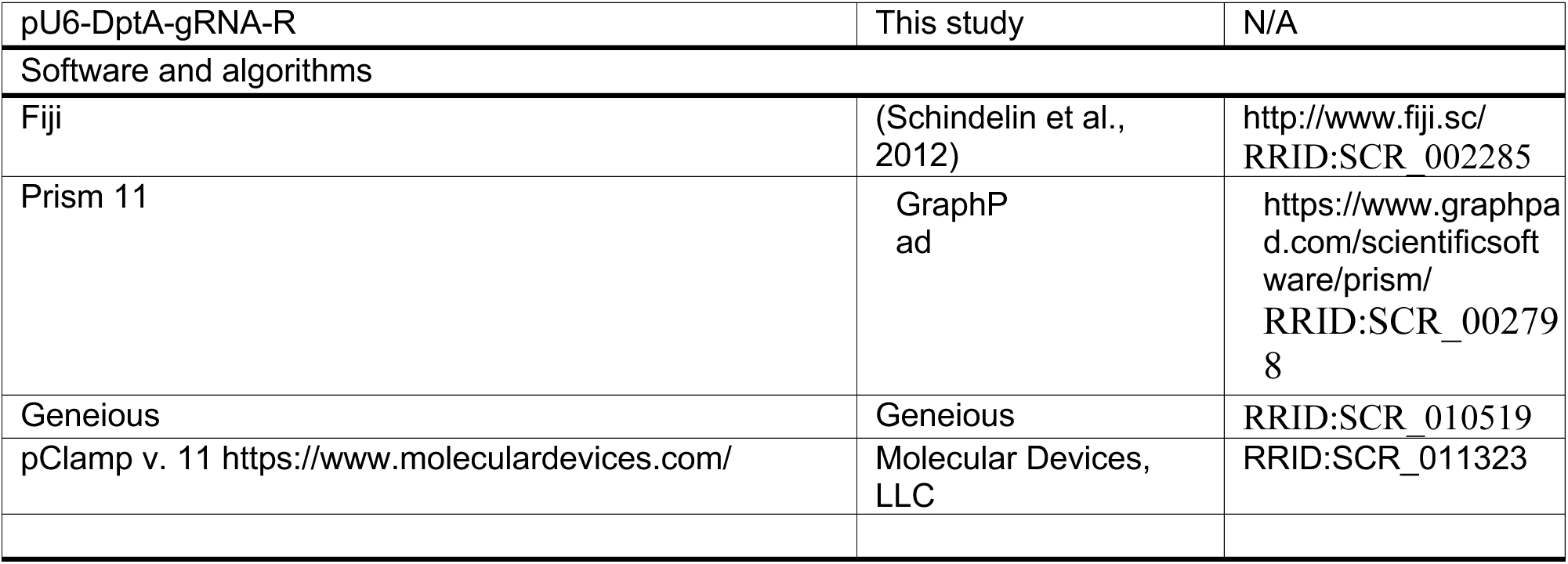

### EXPERIMENTAL MODEL

#### Drosophila stocks and genetics

Flies were raised on Yeast-Cornmeal food (6% cornmeal, 6% yeast, 0.62% agar, 0.1% fruit juice, that was supplemented with 10.6g/L Moldex and 4.9ml/L propionic acid) at 25°C. Experiments were performed on 5-7 days old males at 25°C, unless otherwise stated. Animals were bred and maintained at a low population density in vials and flipped twice a week. Isogenic *w*^1118^ Drosdel flies^86^ (*w^iso^*) were used as wild-type. *Tot^AZX^* and *Tot^XMAZ^* lines have been previously described^48^. Briefly, *Tot^AZX^* flies (lacking TotA, TotB, TotC, TotX and TotZ) were then combined with *TotM* mutation (on the second chromosome) to create the *Tot^XMAZ^* line. U*AS-CecA-HA* was generated by phiC3-mediated recombineering in attP1 site. DptA-HA Knock-In (KI) was generated by CRISPR/cas9-mediated homology-directed repair. A mixture of pScarless-DptA-HA and pU6 vectors containing gRNAs was injected in embryos of the *vasa-cas9* line (BL51324) isogenized in the *w^iso^* background. F1 flies were selected on DsRed eye fluorescence. Stocks used in this study are listed in the Key Resource Table and in **Supplemental Data Table 1**.

### METHOD DETAILS

#### Cloning and DNA constructs

Cloning was performed by In-Fusion Snap Assembly (Takara), following the manufacturer’s instructions. A sequence containing the CecA1 5’UTR and signal peptide was assembled with sequences containing 2 times a HA-tag and CecA1 CDS into the pUASg-attb vector to create the UAS-HA-CecA construct. pScarless-DptA-HA was generated by cloning 1.1Kb upstream of DptA, DptA without its stop codon, a linker encoding 4 times GGS, 3 times a HA-tag, DptA stop codon and 50 bp downstream, a 3xP3-DsRed cassette and 1.2 kb downstream of the DptA coding sequence. The gRNAs used were TATGAGACAATAACCGCCGT and GGGTTAAACAAACAACGCCA were cloned into the pU6 vector by BbsI digestion according to standard protocols.

#### Microbial culture and infection experiments

Bacteria were cultured overnight on a shaking plate at 180 RPM. The following morning, they were pelleted by centrifugation (4000 RPM at 4°C for 15 min) and the bacterial pellets were diluted to an optical density at 600 nm of 200 (OD_600_:200). *Pectobacterium carotovorum carotovorum 15* (*Ecc15*) and *Micrococcus luteus* (*M. luteus*) were grown in LB medium at 29°C. For systemic infections with *Ecc15*, flies were pricked in the thorax with a needle previously dipped into the concentrated bacterial pellet. For experiments with heat killed (HK) bacteria, *Ecc15* and *M. luteus* bacterial pellets diluted to OD_600_:200 were mixed 1:1 and were cyclically (5 times) heated at 95°C and frozen at -20°C. 32 nL of this mixture was injected into the thorax of male adult flies using a nanoinjector (Drummond) and glass capillary needles while control flies were not injected (unchallenged, UC).

#### Negative geotaxis motor ability assay

The climbing ability of flies was measured as previously described^87^. In brief, two empty, transparent Drosophila vials were vertically connected to form a single column, and each vial was marked at its midpoint. For each genotype, 20 adult flies were gently transferred into the apparatus without the use of CO₂ anesthesia to avoid behavioral impairment. Prior to measurement, flies were gently tapped to bring all individuals to the bottom of the column. After 10 s, the positions of the flies within the column were recorded. The column was divided into four equal vertical sections, and a climbing index was assigned based on fly distribution: flies located in the top quarter were assigned a score of 6, those in the second quarter a score of 4, those in the third quarter a score of 2, and those remaining in the bottom quarter a score of 0. The overall climbing index (CI) for each group was calculated following this formula: CI=(N_1_^6+N_2_^4+N_3_^2+N_4_^0)/N_total_, where N₁–N₄ represent the number of flies in each section from top to bottom. Each measurement was performed in triplicate.

#### Electrophysiology

*<u>dSEVC recordings</u>*: were conducted as previously described^54^. Briefly, recordings were conducted in 1.5 mM Ca^2+^ AHL saline. Voltage clamp recordings of adult neuromuscular glutamatergic synaptic currents were carried out using borosilicate glass electrodes (1B120F-4, World Precision Instruments, Florida, USA). Recording electrodes were pulled (Sutter P-1000, California, USA) to resistances of between 20-25 MΩ and filled with 3 M KCl. Suction electrodes (GC120T-10, Harvard Apparatus, Kent, UK) were fire polished to minimize damage to the nerve to a diameter of ∼2–3 µm and filled with extracellular saline. EJCs were evoked (1 ms/0.5 Hz) with a 2100 isolated pulse stimulator (AM Systems, Washington, USA). mEJC and EJC recordings were made from individual muscle fibres of the A2 mvim abdominal muscle segments. Recordings were made using an Axoclamp 900A amplifier (Molecular Devices, Sunnyvale, CA). Muscle fibres were held at −60 mV, recordings sampled at 20 kHz and lowpass filtered at 0.5 kHz using pClamp 11 (Molecular Devices, Sunnyvale, CA). Because Tot-deficient mutants may have altered muscle parameters, muscle inclusion criteria for recordings was more permissive to allow input resistance >5 MΩ rather than >10 MΩ as previously published and injected current <2.5 nA with a view to not exclude meaningful genotype specific physiology. mEJC amplitude and frequency were quantified from 2 minute recordings using automated threshold detection^88^. Only events >100 pA were included to mitigate false positive event detection. EJC amplitudes were averaged from 10 EJCs per recording.

#### Immunostaining and NMJ imaging

Male flies were dissected in PBS along the dorsal midline to expose the abdominal muscles and fixed in 8% paraformaldehyde for 20 minutes at room temperature. The tissues were subsequently rinsed two times with PBS, blocked for 1 hour in PBS containing 1% of goat serum and 2% BSA, and incubated at room temperature for 2 hours in the blocking solution containing a chicken anti-GFP (abcam, 1:1000), mouse anti-HA (Abcam, 1:1000), rat anti-mCherry (Life Technologies, 1:1000) or rabbit anti-TotA^48^ (1:1000) primary antibodies. The tissues were subsequently washed and incubated with an AlexaFluor a488- or a555-coupled secondary antibody (Thermo Fischer Scientific, 1:2000) for 1 hour at room temperature and AlexaFluor a488 anti-HRP (Thermo Fischer Scientific 1:400) to stain motoneurons when stated. After extensive washes, tissues were mounted in glycerol. To assess synaptic bouton morphology, tissues were fixed for 5 min in Bouin’s fixative, permeabilized in PBS-0.1% Triton X-100 for 1 hour, blocked for 1 hour in PBS containing 1% of goat serum and 2% BSA, and incubated overnight at 4°C in the blocking solution containing mouse anti-Brp antibody (nc82, DSHB, 1:100) and chicken anti-Vglut^54^ (1:150) or rat anti-mCherry (Life Technologies, 1:1000) antibodies overnight at 4°C. The tissues were subsequently washed and incubated with appropriate secondary antibodies (Thermo Fischer Scientific, 1:2000) for 1 hour at room temperature. After extensive washes, tissues were mounted in glycerol. Trachea were assessed by imaging the autofluorescent chitin lining the tracheal lumen. The dityrosine bonds chitin fluoresce under UV excitation, allowing chitin to be imaged using a DAPI filter set. Mounted tissues were imaged on a SP8 confocal inverted microscope equipped with 63x oil objective (Leica)

### QUANTIFICATION AND STATISTICAL ANALYSIS

#### Image analysis

Image quantifications were performed using ImageJ software^89^. To quantify AMP and Lactadherin binding at the NMJ, HA or GFP fluorescence intensity was measured on neuronal and glial structures segmented based on HRP-positive signal. Synaptic bouton fragmentation was quantified manually by counting the fraction of motoneuron boutons (vglut- or OK6>mCherry-positive) containing a single active zone (defined as a Brp-positive foci within a bouton). The analysis was performed on motoneurons innervating the *musculi ventralis interni mediales* (mvim) muscles of the A2 segment of male flies from at least 3 independent experiments.

#### Statistical analysis

Each experiment was repeated independently at least three times. Survival curves included experiments with at least one cohort of 20 flies per condition. Survival analyses were performed using a log-rank test (Mantel-Cox). Quantification data were analyzed using a Mann-Whitney test or ordinary one-way ANOVA with Dunnett’s multiple comparisons test, as stated in the figure legends. Each dot represents an individual NMJ, horizontal bars represent the median. Error bars represent the standard deviation of the mean of replicate experiments (SD). *p*-values are represented in the figures by the following symbols: ns for p ≥ 0.05, ^∗^ for *P* between 0.01 and 0.05; ^∗∗^ for *P* between 0.001 and 0.01, ^∗∗∗^ for *P* between 0.0001 and 0.001, ^∗∗∗∗^ for p ≤ 0.0001. Multiple comparison statistics were represented using a compact letter display graphical technique: groups were assigned the same letter if they were not significantly different (P > 0.05).

