## Supplementary figures and images for "The phosphatidylserine-binding proteins Turandots protect the peripheral nervous system from antimicrobial peptide toxicity"

### Figure S1

A

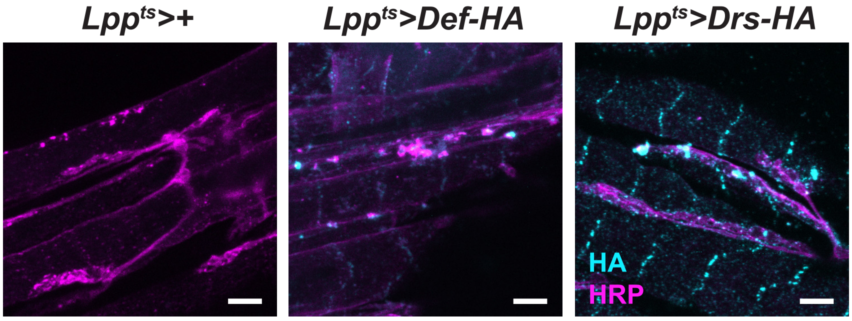

### Figure S2

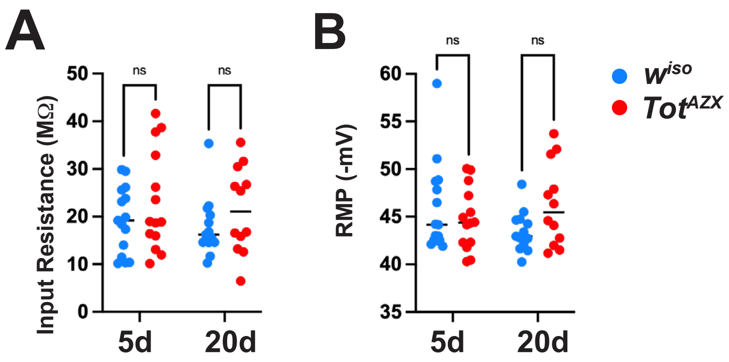
