## Supplemental Table S1 for "The phosphatidylserine-binding proteins Turandots protect the peripheral nervous system from antimicrobial peptide toxicity"

| | **name** | **genotype** | **origin** |
|---|---|---|---|
| | dptA-HA KI | wiso;dptA-HA;+ | this study |
| | UAS-DptA-HA | +;;UAS-DptA-HA | FlyORF F002330 |
| | *$Lpp^{ts}$ >* | *w;tub-gal80$^{ts}$;lpp-gal4/TM3sb* | Brankatschk and Eaton 2010 |
| | Rel IR | w;Rel IR | VDRC KK |
| Figure 1 | *UAS-lactGFP* | *w;UAS-GFP-LactC1C2/CyO-GFP* | Sapar et al. 2018 |
| | vglut::Tdt | Vglut1-LexA, 13xLexAop2-CD4-TdTomato/CyO | BDSC 60314 and 77138 |
| | vglut::Tdt; Lpp> | Vglut1-LexA, 13xLexAop2-CD4-TdTomato/CyO; Lpp-gal4/TM3sb | BDSC 60314 and 77138 and Brankatschk and Eaton 2010 |
| | LexO-xkr8,UAS-dptA-HA | LexAop-xkr8, UAS-dptA-HA/TM3sb | Sapar et al. 2018 and FlyORF F002330 |
| | *DptA-HA,UAS-scramb1 IR* | *;UAS-scramb1 IR, dptA-HA* | VDRC 107024 and this study |
| | OK6>mCherry | *OK6-gal4, UAS-6xmCherry/CyO* | Banerjee et al. 2021 |
| | *OK6>mCherry $Tot^{AZX}$* | *;OK6-gal4, UAS-6xmCherry;TotX$^{JP44}$, TotAZ* | This study |
| | dptA-HA KI | wiso;dptA-HA;+ | this study |
| | *wiso;dptA-HA;$Tot^{AZX}$* | wiso;dptA-HA;+ | this study |
| | *Lpp>* | *w;;lpp-gal4/TM3sb* | Brankatschk and Eaton 2010 |
| Figure 2 | UAS-CeA-HA | ;;UAS-CecA-HA | this study |
| | *UAS-CecA-HA;$Tot^{AZX}$* | *;UAS-CecA-HA;$Tot^{AZX}$* | this study |
| | *$w^{iso}$* | *iso $w^{1118}$ DrosDel* | Ferreira et al. 2014 |
| | *$Tot^{AZX}$* | *iso;iso;TotX$^{JP44}$, TotAZ* | Rommelaere et al. 2024 |
| | *AMP12,$Tot^{AZX}$* | *iso;Def$^{SK3}$, AttC$^{Mi}$, Dro-AttAB$^{SK2}$, Mtk$^{R1}$, Dpt$^{SK1}$; iso;iso;CecKOΔA-C,TotX$^{JP44}$, TotAZ$^{SK6}$* | Rommelaere et al. 2024 |
| | *CecAC,$Tot^{AZX}$* | *iso;iso;CecKOΔA-C,TotX$^{JP44}$, TotAZ$^{SK6}$* | Rommelaere et al. 2024 |
| | *UAS-PGRP-LC* | *w;; UAS-PGRP-LCx-FL* | Neyen et al. 2017 |
| | *$Lpp^{ts}$ >* | *w;tub-gal80$^{ts}$;lpp-gal4/TM3sb* | Brankatschk and Eaton 2010 |
| | UAS-DptA-HA | +;;UAS-DptA-HA | FlyORF F002330 |
| | *UAS-Def-HA* | *;;UAS-Def.ORF.3xHA.GW @attP86Fb* | FlyORF: F002467 |
| | *UAS-Drs-HA* | *;;UAS-Drs.ORF.3xHA.GW @attP86Fb* | FlyORF: F002219 |
| | *$w^{iso}$* | *iso $w^{1118}$ DrosDel* | Ferreira et al. 2014 |
| Figure 3 | *$Tot^{AZX}$* | *iso;iso;TotX$^{JP44}$, TotAZ* | Rommelaere et al. 2014 |
| | *$Tot^{XMAZ}$* | *iso;totM$^{JP1621}$;TotX$^{JP44}$, TotAZ$^{SK6}$* | Rommelaere et al. 2014 |
| | OK6>mCherry | *OK6-gal4, UAS-6xmCherry/CyO* | Banerjee et al. 2021 |
| | *OK6>mCherry $Tot^{AZX}$* | *;OK6-gal4, UAS-6xmCherry;TotX$^{JP44}$, TotAZ* | This study |
| | *OK6>mCherry $Tot^{XMAZ}$* | *;OK6-gal4, UAS-6xmCherry, totM$^{JP1621}$;TotX$^{JP44}$, TotAZ$^{SK6}$* | This study |
| | *UAS-trio;$Tot^{AZX}$* | *;;UAS-trio, TotX$^{JP44}$, TotAZ$^{SK6}$* | BDSC 9513 and this study |
| | *AMP12,$Tot^{AZX}$* | *iso;Def$^{SK3}$, AttC$^{Mi}$, Dro-AttAB$^{SK2}$, Mtk$^{R1}$, Dpt$^{SK1}$; iso;iso;CecKOΔA-C,TotX$^{JP44}$, TotAZ$^{SK6}$* | Rommelaere et al. 2024 |
| | *$Lpp^{ts}$ >* | *w;tub-gal80$^{ts}$;lpp-gal4/TM3sb* | Brankatschk and Eaton 2010 |
| | *UAS-lactGFP* | *w;UAS-GFP-LactC1C2/CyO-GFP* | Sapar et al. 2018 |
| | *OK6>mCherry $Tot^{AZX}$* | *;OK6-gal4, UAS-6xmCherry;TotX$^{JP44}$, TotAZ* | This study |
| | *UAS-ATP8 $Tot^{AZX}$* | *;UAS-CDC50-T2A-ATP8A; TotX$^{JP44}$, TotAZ$^{SK6}$* | Ji et al. 2023 and this study |
| | *UAS-scramb1 IR $Tot^{AZX}$* | *;UAS-scramb1 IR; TotX$^{JP44}$, TotAZ$^{SK6}$* | Rommelaere et al 2024 |
| | *MHC>; $Tot^{AZX}$* | *;MHC-RFP,MHC-gal4;TotX$^{JP44}$, TotAZ* | BDSC 38464 and this study |
| Figure 4 | OK6>mCherry | *OK6-gal4, UAS-6xmCherry/CyO* | Banerjee et al. 2021 |
| | *MHC>* | *;MHC-RFP,MHC-gal4* | BDSC 38464 |
| | *UAS-xkr8* | *;UAS-xkr8/CyoGFP* | Sapar et al. 2018 |
| | *OK6>mCherry; $CecAC^{\Delta}$* | *;OK6-gal4, UAS-mCherry; CecKOΔA-C* | Banerjee et al. 2021 and Carboni et al. 2022 |
| | *UAS-xkr8; $CecAC^{\Delta}$* | *;UAS-xkr8/CyoGFP; CecKOΔA-C* | Rommelaere et al. 2024 |
| | *UAS-drpr IR;$Tot^{AZX}$* | *;UAS-drpr IR; TotX$^{JP44}$, TotAZ$^{SK6}$* | BDSC 67034 and this study |
| | *UAS-SIMU IR;$Tot^{AZX}$* | *;UAS-SIMU IR; TotX$^{JP44}$, TotAZ$^{SK6}$* | BDSC 61866 and this study |
